# The Transcriptional landscape of *Streptococcus pneumoniae* TIGR4 reveals a complex operon architecture and abundant riboregulation critical for growth and virulence

**DOI:** 10.1101/286344

**Authors:** Indu Warrier, Nikhil Ram-Mohan, Zeyu Zhu, Ariana Hazery, Haley Echlin, Jason Rosch, Michelle M Meyer, Tim van Opijnen

## Abstract

Efficient and highly organized regulation of transcription is fundamental to an organism’s ability to survive, proliferate, and quickly respond to its environment. Therefore, precise mapping of transcriptional units and understanding their regulation is crucial to determining how pathogenic bacteria cause disease and how they may be inhibited. In this study, we map the transcriptional landscape of the bacterial pathogen *Streptococcus pneumoniae* TIGR4 by applying a combination of high-throughput RNA-sequencing techniques. We successfully map 1864 high confidence transcription termination sites (TTSs), 790 high confidence transcription start sites (TSSs) (742 primary, and 48 secondary), and 1360 low confidence TSSs (74 secondary and 1286 primary) to yield a total of 2150 TSSs. Furthermore, our study reveals a complex transcriptome wherein environment-respondent alternate transcriptional units are observed within operons stemming from internal TSSs and TTSs. Additionally, we identify many putative *cis*-regulatory RNA elements and riboswitches within 5’-untranslated regions (5’-UTR). By integrating TSSs and TTSs with independently collected RNA-Seq datasets from a variety of conditions, we establish the response of these regulators to changes in growth conditions and validate several of them. Furthermore, to demonstrate the importance of ribo-regulation by 5’-UTR elements for *in vivo* virulence, we show that the pyrR regulatory element is essential for survival, successful colonization and infection in mice suggesting that such RNA elements are potential drug targets. Importantly, we show that our approach of combining high-throughput sequencing with *in vivo* experiments can reconstruct a global understanding of regulation, but also pave the way for discovery of compounds that target (ribo-)regulators to mitigate virulence and antibiotic resistance.

**Author summary:** The canonical relationship between a bacterial operon and the mRNA transcript produced from the operon has become significantly more complex as numerous regulatory mechanisms that impact the stability, translational efficiency, and early termination rates for mRNA transcripts have been described. With the rise of antibiotic resistance, these mechanisms offer new potential targets for antibiotic development. In this study we used a combination of high-throughput sequencing technologies to assess genome-wide transcription start and stop sites, as well as determine condition specific global transcription patterns in the human pathogen *Streptococcus pneumoniae*. We find that the majority of multi-gene operons have alternative start and stop sites enabling condition specific regulation of genes within the same operon. Furthermore, we identified many putative RNA regulators that are widespread in the *S. pneumoniae* pan-genome. Finally, we show that separately collected RNA-Seq data enables identification of conditional triggers for regulatory RNAs, and experimentally demonstrate that our approach may be used to identify drug-able RNA targets by establishing that pyrR RNA functionality is critical for successful *S. pneumoniae* mouse colonization and infection. Thus, our study not only uses genome-wide high-throughput approaches to identify putative RNA regulators, but also establishes the importance of such regulators in *S. pneumoniae* virulence.

## Introduction

The transcriptional architecture of bacterial genomes is far more complex than originally proposed. The classical model of an operon describes a group of genes under the control of a regulatory protein where transcription results in a polycistronic mRNA with a single transcription start site (TSS) and a single transcription terminator site (TTS) [1]. However, many individual examples have established that the same operon may encode alternative transcriptional units under varying environmental conditions [2,3]. Furthermore, advancements in sequencing technology that enable highly accurate mapping of TSSs and TTSs on a genome-wide level have demonstrated that the number of TSSs and TTSs can significantly exceed the number of operons [4]. Thus it seems likely that the bacterial transcriptional landscape, or the genome-wide map of all possible transcriptional units, is shaped by an operon architecture that encodes many TSSs and TTSs within single operons, thus significantly increasing complexity with the objective of enabling diverse transcriptional outcomes [5,6].

To achieve a complex landscape of alternative transcriptional units, transcriptional regulation occurs on multiple levels. In addition to the many protein activators and repressors that control transcription initiation, there are also many non-coding RNAs (ncRNAs), including both small ncRNAs (sRNAs) and highly structured portions of mRNAs that play essential roles as regulatory elements controlling metabolism, stress-responses, and virulence [7-9]. *Trans –* acting small RNAs (sRNAs), which are found in intergenic regions [10] or derived from 3’ UTRs [11], allow selective degradation or translation of specific mRNAs [10]. *Cis –* acting mRNA structures, such as riboswitches, which interact with small molecules including metal ions, and protein ligands, and other regulatory sequences that are found in the long 3’ UTR of mRNAs, affect expression of their respective genes by regulating transcription attenuation or translation inhibition [12,13]. RNA regulation has been shown to play a key role in shaping the transcriptional landscape of a wide range of pathogenic bacteria including *Staphylococcus aureus*, *Listeria monocytogenes, Helicobacter pylori,* and strains of *Streptococci* [6, 14-22]. Several RNA regulators have been validated and associated with pathogenicity and virulence [23,24], and could be used as highly specific druggable targets [25,26], however, only a select set of regulators have been targeted to date [27,28].

*Streptococcus pneumoniae* is a major causative agent of otitis media, meningitis, pneumonia, and bacteremia. It causes 1.2 million cases of drug-resistant infection in the US annually and results in ~1 million deaths per year worldwide [29-31]. While high-resolution transcriptional mapping data are available for other *Streptococcus* species, these studies have shown limited experimental validation [20], or have focused primarily on the role of sRNAs in virulence [32]. Additionally, previous studies of the *S. pneumoniae* transcriptome have demonstrated the presence of ncRNA regulators and assessed their roles in infection and competence, however, these studies also largely focused on sRNAs [16,18,33]. Thus high-resolution validated maps of the genome wide transcriptional landscape for different *S. pneumoniae* strains would be incredibly valuable.

Here a comprehensive characterization of the *S. pneumoniae* TIGR4 transcriptional landscape is created using single and paired-end RNA-Seq [34], 5’ end-Seq [22], and term-seq (3’-end sequencing) [35] during early-log and mid-log growth in the presence and absence of the antibiotic vancomycin. We obtain a global transcript coverage map, identifying all TSSs, and all TTSs, which highlights a highly complex *S. pneumoniae* transcriptional landscape including many operons with multiple TSSs and TTSs. Furthermore, we demonstrate how TSS and TTS mapping under one set of conditions can be leveraged to analyze independently obtained RNA-Seq data collected under a variety of conditions, and we experimentally validate this approach with several cis-acting RNA regulators. Finally, we demonstrate that the functionality provided by the RNA *cis* – regulator *pyrR* is critical for *S. pneumoniae in vivo* in a mouse infection model. Importantly, our work demonstrates how a variety of high-throughput sequencing efforts can be combined to map out a comprehensive transcriptional landscape for a bacterial pathogen as well as identify potentially druggable ncRNA targets.

## Results and Discussion

### Streptococcus pneumoniae has a complex transcriptional landscape

To characterize the transcriptional landscape of *Streptococcus pneumoniae* TIGR4 (T4), we first determined transcript boundaries by mapping transcription start (TSSs) and termination sites (TTSs) from 5’ and 3’ end sequencing reads obtained from 3 different conditions: early-log and mid-log growth phase in the absence and presence of vancomycin (Fig 1A and 1B). Since the TSS predictions (both whether a TSS exists and the position identified) correspond well among all three conditions for the vast majority of genes (S1 Fig), reads from both mid-log phase growth conditions were pooled for downstream analyses. To describe a comprehensive transcriptional landscape for T4, the 2341 annotated genes were supplemented by annotating the homologs of known structured non-coding RNAs (ncRNAs) in T4 identified using Infernal (an RNA specific homology search tool [36]), and previously described small RNAs (sRNAs) [16,18] resulting in a total of 2557 features in T4. Given this feature set, a total of 742 primary, and 48 secondary, high confidence TSSs were identified, with a processed coverage greater than 100 and a processed/unprocessed ratio greater than 4, where the primary TSS has the highest processed/unprocessed ratio. A total of 1286 low confidence primary TSSs were called that have coverage greater than 2 and a processed/unprocessed ratio greater than 1. In addition, 74 low confidence secondary TSSs that have coverage greater than 100, but a lower processed/unprocessed ratio than the primary TSS were called to yield 2150 total TSSs. From the pooled 3’-end sequencing reads, 1864 TTSs were identified that have a minimum coverage of 10, and are enriched at least two-fold over the background (Fig 2, S1 and S2 Tables). Of the 742 high confidence TSSs, 625 showed the presence of the GRTATAAT motif at the −10 position and 215 had the MTTGAMAA motif at the −35 position (S1 Fig).

**Figure 1.**
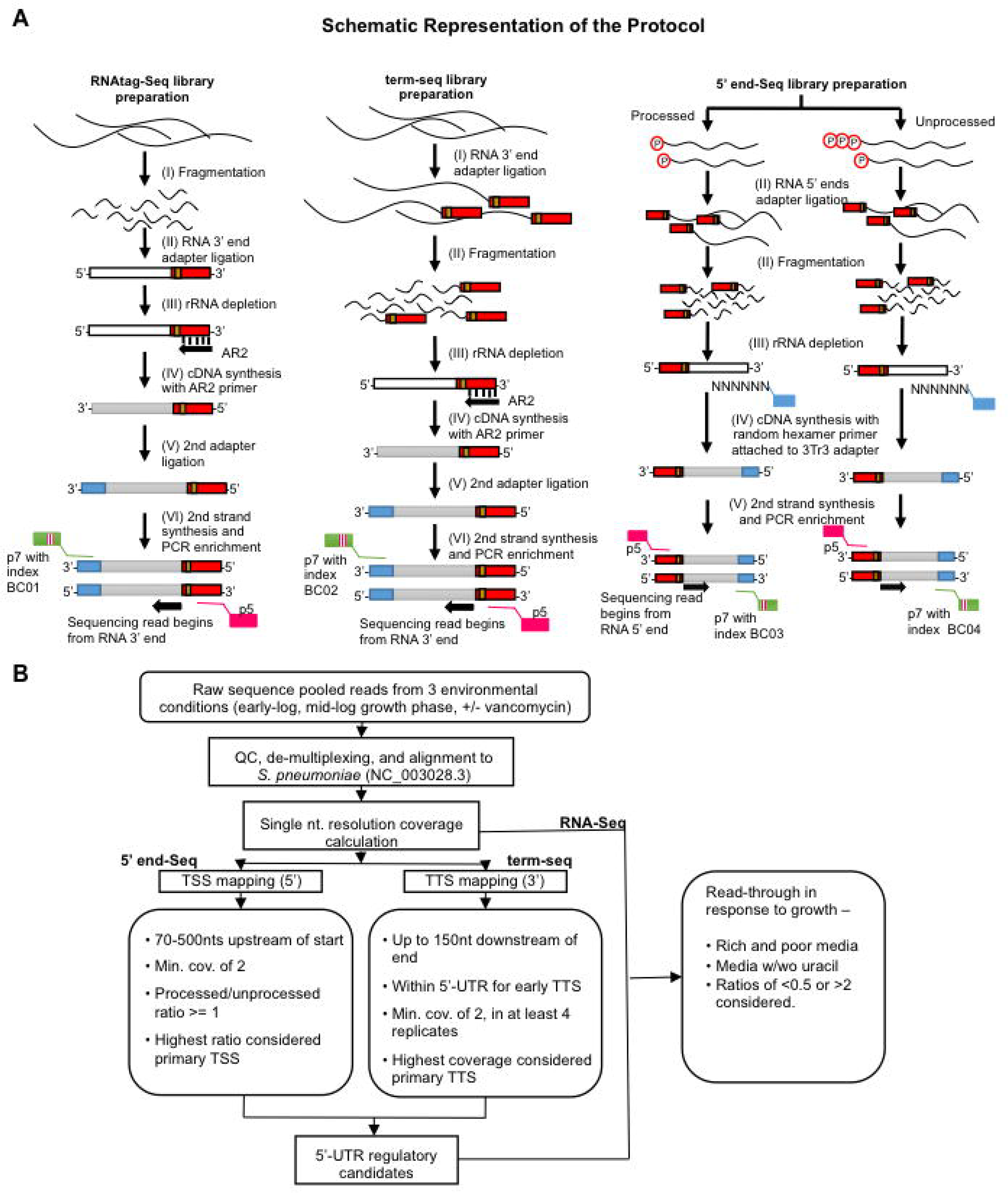
Schematic representation of the sequencing and data analysis methodology. **(A)** A description of the experimental pipeline. RNA-Seq (left column) and term-seq (middle column) libraries were prepared according to protocols described in [34,35]. 5’ end-Seq libraries (right column) were generated by dividing the total RNA into 5’ polyphosphate treated (processed) and untreated (unprocessed) samples and subsequently processed according to protocols described in [22,64]. White and grey lines correspond to RNA and cDNA, respectively and colored blocks represent unique sequence barcodes. Illumina sequencing adaptors with index barcodes are also indicated. **(B)** A brief description of the analysis pipeline. Raw reads from all three sequencing methodologies obtained from three environmental conditions (early-log and mid-log growth phase in the presence and absence of vancomycin) were de-multiplexed and aligned to T4 (NC_003028.3). Based on the reads mapped, single nucleotide coverage was calculated. Coverage of the 5’ end of the reads calculated for the 5’ end-Seq was used to determine the transcription start sites. Coverage of the 5’ end of the reads from term-seq was used to determine the transcription termination sites. RNA-Seq coverage was used to calculate the read-through across candidate 5’ untranslated regions with early transcription terminators. Read-through responses of the candidates in different media conditions (rich and poor media, media with and without uracil) were analyzed to identify environment-responsive RNA regulators.

**Figure 2.**
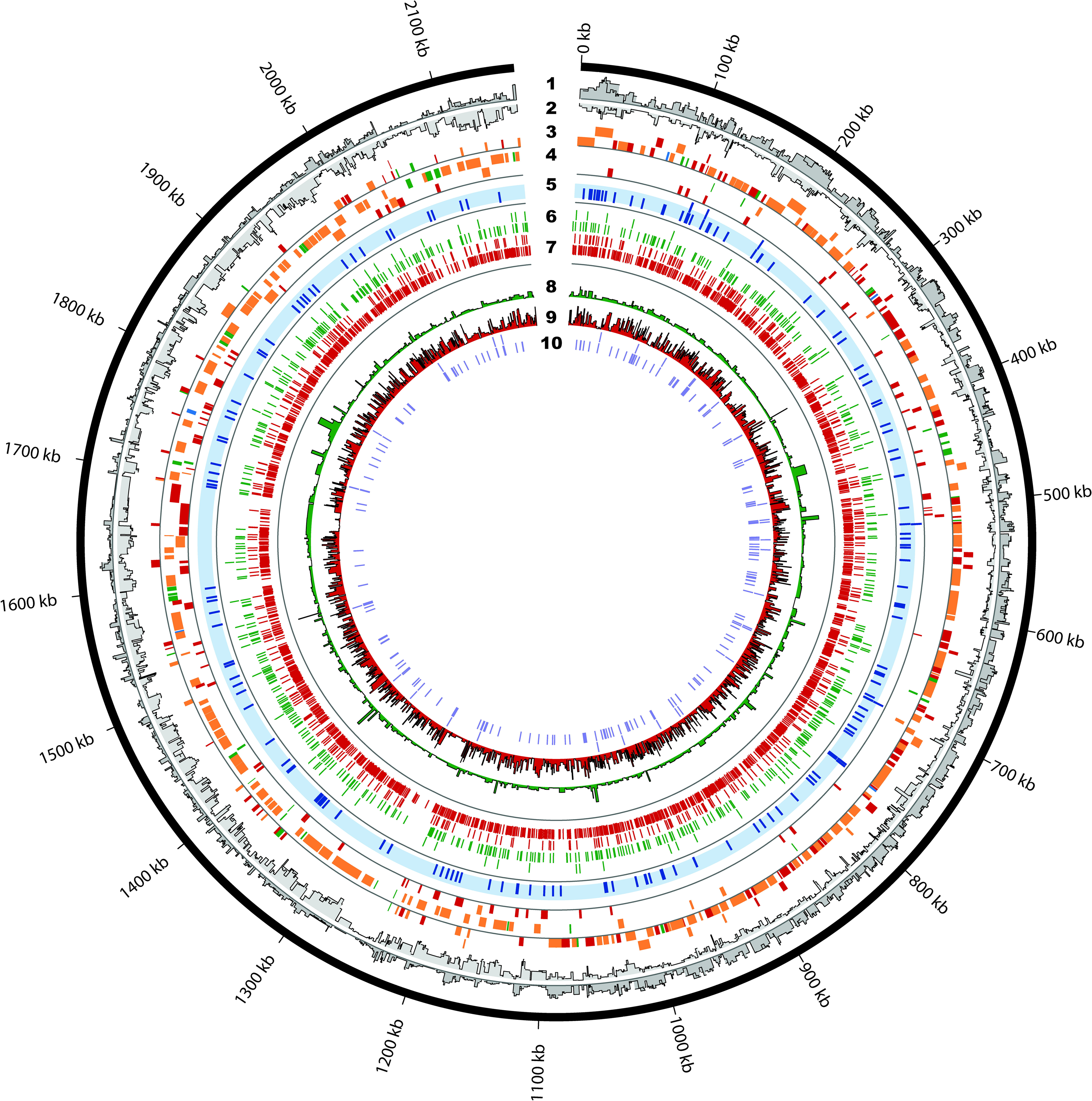
Genome-wide map of the *Streptococcus pneumoniae* TIGR4 transcriptional landscape. A map of all the transcriptional features identified. **1.** Paired-end RNA-Seq coverage of the plus strand. **2.** Paired-end RNA-Seq coverage of the minus strand. **3.** Annotated operon structures on the plus strand. Tile colors represent classification of the operons according to their number of high confidence TSS and TTSs: 1) traditional operons consisting of multiple genes with a single TSS and a TTS (green); 2) multiTSS operons consisting of multiple genes with internal TSSs but one TTS (blue); 3) multiTTS operons consisting of multiple genes with a single TSS but multiple internal TTSs (red); and 4) complex operons consisting of multiple genes with multiple internal TSSs and TTSs (orange). To avoid clutter, simple operons consisting of a single gene with a single TTS and TSS are omitted. **4.** Annotated operon structures on the minus strand. **5.** 141 novel putative intergenic regulatory elements in blue. **6.** 742 high confidence transcriptional start sites (TSS) in green. **7.** 1864 enriched transcriptional termination sites (TTS) in red. **8.** Log transformed processed/unprocessed ratio of each high confidence TSS. **9.** Log transformed coverage of each predicted TTS. **10.** Positions of the known structured non-coding RNAs and small RNAs. Figure was generated using Circos [79].

In order to assess how the high confidence TSSs and TTSs contribute to transcriptional complexity, we reconstructed T4 transcripts from paired-end RNA-Seq data. StringTie [37,38] detected 343 single gene operons and grouped 1857 genes into 388 multi-gene operons (S3 Table, and Fig 3). Based on the reconstructed transcripts, we classified operons into five categories based on the number of internal primary high confidence TSSs and TTSs (Fig 2). Simple operons, transcriptional units with a single gene TSS, and TTS are predominantly identified in the *S. pneumoniae* genome at 47% of all operons (Fig 3A). Traditional operons with multiple genes and a single TSS and TTS make up 10% of the operons, while multiTSS operons make up 1%, and multiTTS operons make up 16%, of all transcriptional units respectively. However, complex operons (most of which consist of two genes) with multiple TSSs and TTSs are the second largest category comprising 26% of all operons (Fig 3A and Fig 3B). Most complex operons are defined by a secondary internal TSS and TTS, however there are several significantly more complex examples where the operon contains multiple TSSs and TTSs (Fig 3B), indicating an intricate system of possible transcripts. Of note, transcript prediction from paired-end RNA-Seq using StringTie agrees partially with operon predictions run using only single-end RNA-Seq data [39], or established operon databases that use genome context [40]. Such discrepancies between strategies for operon calling have been observed previously (e.g. *Escherichia coli* MG1655) [4,41]). For example, StringTie groups genes SP_0001-SP_0014 into a single transcript while Rockhopper splits them into three operons. However, it is evident from the paired-end RNA-Seq coverage map (see S2 Fig) that despite the fluctuations in coverage between the coding regions, there is still large enough expression in the intergenic regions for StringTie to group these genes together. This example demonstrates the role of internal transcriptional features in creating alternative transcriptional units. Of note, operon predictions based on either method can only be validated by directed, low-throughput approaches. However, despite the differences arising from the methodologies used, the complexity in operon structure we deduce remains nearly the same except for the portion of multiTTS operons (see S2 Fig). The TSS, TTS, our supplemented annotations, and reconstructed transcripts can be easily visualized using files available at https://github.com/nikhilram/T4pipeline.

**Figure 3.**
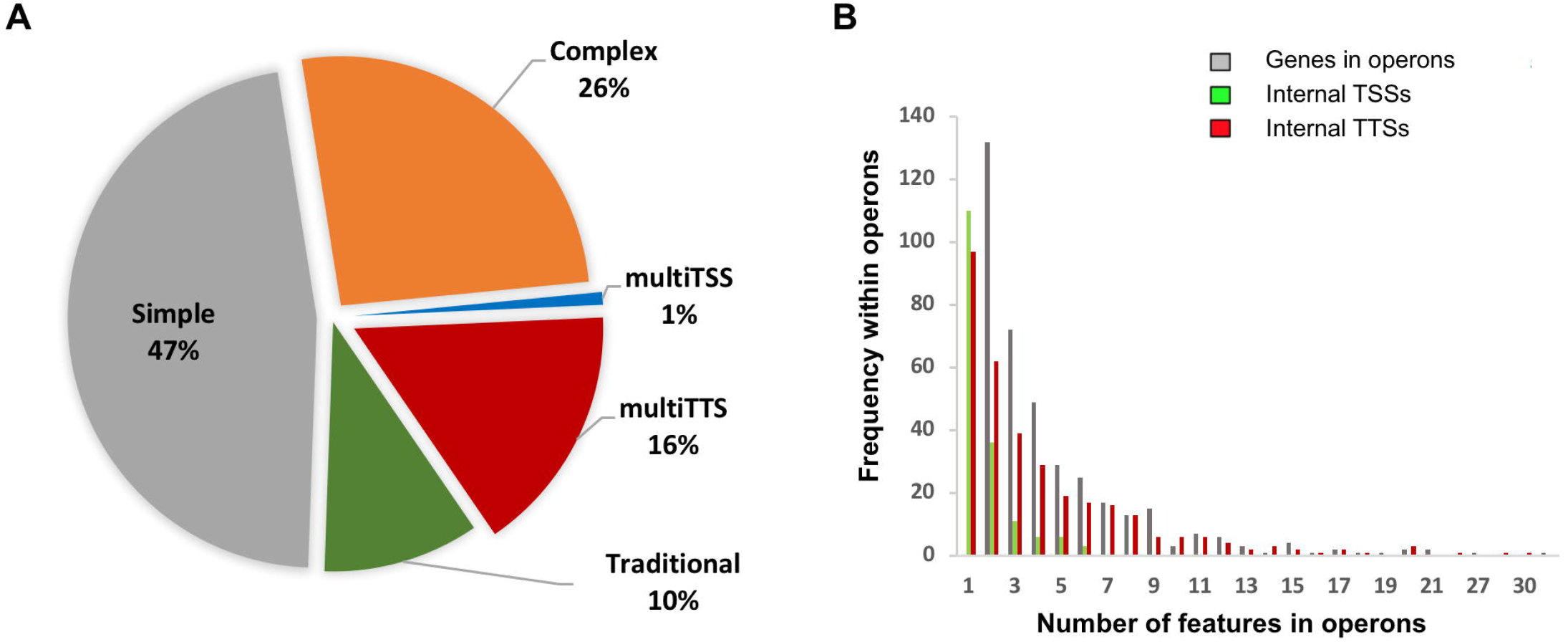
Distribution of operon types in the genome and frequency of transcriptional features within non-traditional operons identified using StringTie. **(A)** The pie chart describes the distribution of the types of operons present in T4 annotated using only high confidence TSSs. A total of 388 multigene and 343 single gene operons were identified, which can be divided up in 47% simple operons (single gene transcriptional units with a single TSS and TTS; gray), 26% complex operons (multi-gene operons with multiple TSSs and TTSs; orange), 10% traditional operons (multi-gene operon with a single TSS and TTS; green), 1% multiTSS operons (blue), and 16% multiTTS operons (red). (**B)** The clustered histogram describes the distribution of genes and transcriptional features in non-traditional operons, where gray represents the numbers of genes in the multigene operons, green represents the number of TSSs within operons, red represents the number of TTSs within operons. Two-gene operons are found most frequently in the non-traditional operons with one internal TSS and TTS.

Since our data revealed many operons with complex structure, we sought to corroborate specific examples using additional data sources. One complex operon we identified, which is also present in existing databases of operon structure [5,42], consists of 9 genes (SP_1018-SP_1026) encoding thymidine kinase, GNAT family N-acetyltransferase, peptide chain release factor 1, peptide chain release factor N(5)-glutamine methyltransferase, threonylcarbomyl-AMP synthase, N-acetyltransferase, serine hydroxymethyltransferase, nucleoid-associated protein, and Pneumococcal vaccine antigen A respectively, with two primary high confidence and six other internal TSSs and eight TTSs (Fig 4A). In addition, this operon displays unequal and complex gene expression patterns when independently collected RNA-Seq data from diverse media conditions is mapped to the transcript. In poor growth medium (MCDM) the operon can be split into two parts based on expression observed from RNA-Seq coverage maps, where the last five genes in the operon (SP_1022-1026) are expressed higher than the first four genes (SP_1018-1021), while in rich medium (SDMM) the read depth across the operon is similar. While none of the individual genes show significant differential expression between the two conditions (>2 fold change [43,44], there is a drastic difference between the two conditions in the average coverage per gene amongst the first 4 versus the last 5 genes. In rich medium, SP_1022-1026 show an average coverage of ~1.9x more than SP_1018-1021, while in the poor medium the average coverage of SP_1022-1026 is ~3.6x that of SP_1018-1021. This observation corroborates the role of the internal regulatory mechanisms to create alternate transcriptional units for maintaining differences in gene expression between different growth conditions especially since we identify a high confidence TSS upstream of SP_1022.

**Figure 4.**
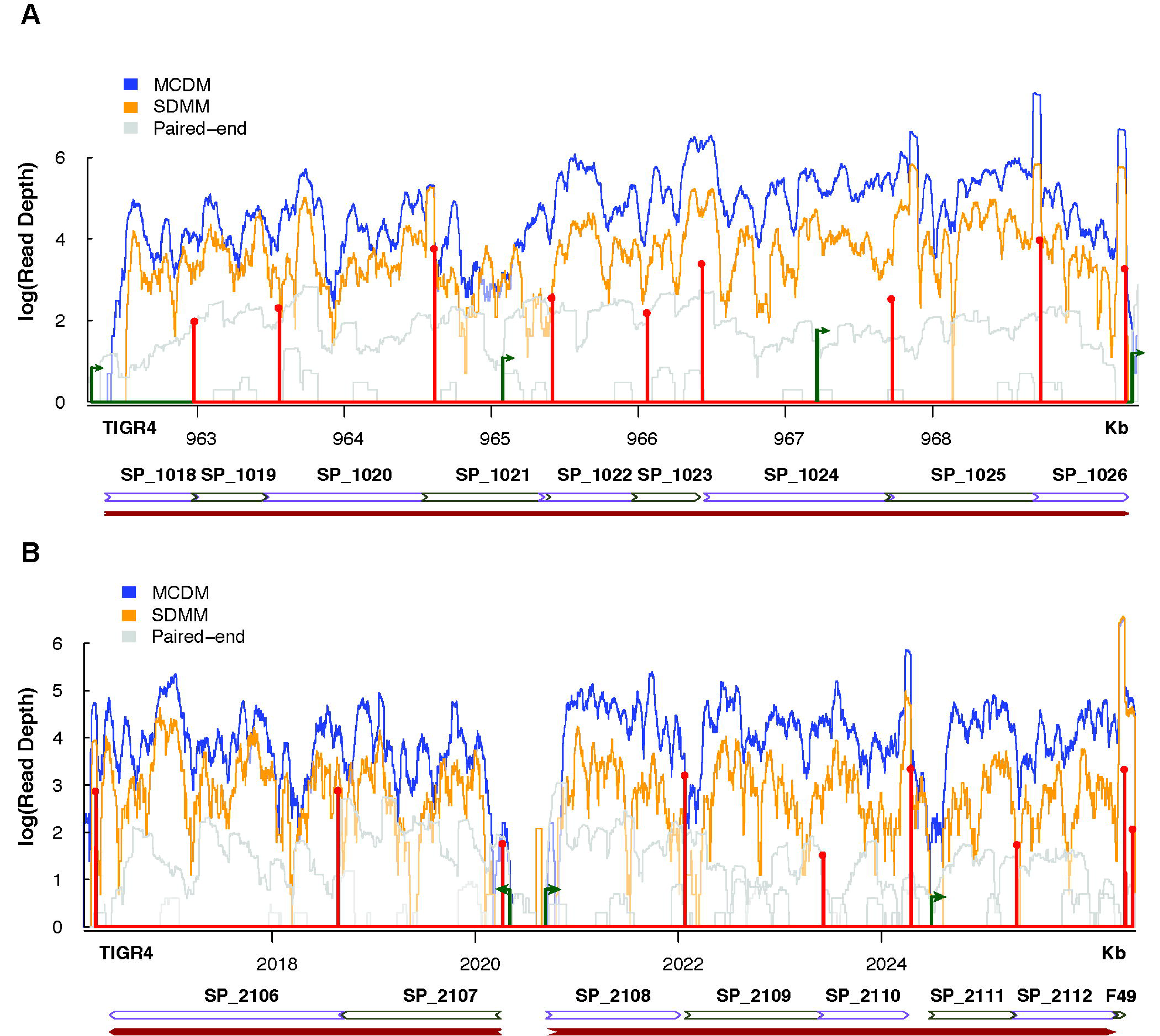
Variability in expression levels between genes in the same operon when grown in rich (SDMM) and poor (MCDM) media conditions. RNA-Seq coverage maps of complex operon/regulon including the TSSs in green and TTSs in red. Size of the transcriptional features represents log transformation of the processed/unprocessed ratio for TSSs and coverage for TTSs. Arrows on the TSS represent the strand and direction of transcription. There are little to no reads mapped to the strand opposite to coding regions. Coverage was normalized to the number of uniquely mapped reads in each library and log_10_ transformed for representation (**A)** A complex 9-gene operon (SP_1018–SP_1026) encoding thymidine kinase, GNAT family N-acetyltransferase, peptide chain release factor 1, peptide chain release factor N(5)-glutamine methyltransferase, threonylcarbomyl-AMP synthase, N-acetyltransferase, serine hydroxymethyltransferase, nucleoid-associated protein, and Pneumococcal vaccine antigen A respectively. Genes SP_1022-SP_1026 are expressed to greater levels in MCDM than in SDMM unlike genes SP_1018-SP_1021. (**B)** The maltose regulon, an example of two operons of different complexities working together in response to maltose in the medium. Complex operon SP_2108-SP_2112 shows greater expression in MCDM than SDMM in comparison to the multiTTS operon SP_2106-SP_2017

Additional validation of our data and analysis approach derives largely from existing low-throughput experiments. The *mal* regulon is a multiple operon system under the control of a single protein, *malR* (SP_2112), which downregulates regulon expression at the *malM* (SP_2107) promoter [45]. Our data shows that the mal regulon includes operons belonging to two different categories, a multiTTS operon (*malMP*/SP_2016-2107) and a complex operon (*malXCD*/SP_2108-2112). From the RNA-Seq coverage maps it is clear that the two operons can be differentially controlled and expressed in rich vs poor media (Fig 4B). Again, although the genes are not significantly differentially expressed in the two conditions [43,44], the average coverage for each gene is greater in the poor medium in comparison to that in the rich medium.

Furthermore, the TSS and TTS identified by our analysis reveal features that have been previously described in lower throughput assays [24]. Despite experimental analyses grouping SP_2108-2110 and SP_2111-2112 into two different operons, it is evident from the paired-end RNA-Seq coverage map as well as RNA-Seq of T4 grown in MCDM that transcription continues past the TTS of SP_2110 into SP_2111. In SDMM, however, transcription seems to mimic what has been experimentally shown. Thus, although our data may highlight many examples of complex transcriptional architecture, these examples are verifiable through the incorporation of additional RNA-Seq data, and where applicable are consistent with low-throughput studies done in the past.

### Genome-wide identification and pan-genome wide conservation of regulatory RNAs

To identify RNA regulators that act through premature transcription termination, we compiled 3’ end sequencing reads upstream of translational start sites, allowing a minimum 5’-untranslated region (UTR) length of 70 bases. We detected 162 such early TTS sites that represent novel high and low confidence putative regulatory elements, of which 141 were completely intergenic (represented in blue in the 5th band of Fig 2). Intriguingly, these putative regulatory elements are not limited to the 5’-UTR of protein coding genes. Of these 141 novel putative regulatory elements, 129 are upstream of protein coding genes while the rest are in the 5’-UTR of annotated ncRNAs. A ncRNA under the regulation of another ncRNA has been shown in *Enterococcus faecalis* [46] and *Listeria monocytogenes* [47] but this is the first putative example of this mechanism in *Streptococcus pneumoniae* TIGR4. By screening these regulatory elements against 380 published *S. pneumoniae* strains [48], covering a large part of the pan-genome, we found that 80 candidates (~57%) were conserved across all genomes, 107 (76%) were identified in at least 350 genomes, 12 (~9%) candidates were identified in fewer than half of the genomes (S3A Fig), while the least distributed candidate, identified upstream of a hypothetical protein, was found only in 52 genomes analyzed. Interestingly, 140 (99%) candidates were found as single copies within a genome, while copies of the putative candidate upstream of the large subunit of the rRNA were found preceding each of the 3 rRNA copies in each genome. Evolutionary distance of each candidate cluster was estimated using MEGA-CC [49], which reveals that each cluster is made of highly similar, if not identical, sequences (S3B Fig).

### Leveraging RNA-Seq data collected under various conditions enables identification and validation of environment-responsive RNA regulators

We reasoned that since we mapped RNA-regulators by means of 5’end and 3’end sequencing, we would be able to associate these regulators with specific growth conditions using environment dependent RNA-Seq data. To confirm the biological relevance of an RNA regulator and associate it with a specific condition one would expect to see a change in the 5’ UTR coverage relative to the accompanying gene. For instance, if a regulator forms an early terminator the RNA-Seq coverage in the 5’UTR is relatively high, while the coverage in the controlled gene would be much lower. Alternatively, if the environment relieves the formation of the early terminator the coverage across the 5’UTR and gene would become less skewed. To determine the applicability of this assumption we leveraged independently collected RNA-Seq data sampled under different nutrient conditions including, rich and poor media [44], and nutrient depletion conditions where a single nutrient was removed from the environment. RNA-Seq data were mapped to each putative regulatory region and coverage was calculated and averaged across the length of the 5’ UTR regulatory element and the downstream gene. From our list of candidate 5’-UTR regulatory elements, 36 showed more than two fold change in read-through between rich and poor media, with the majority showing an increase in readthrough in poor media. Furthermore, as more RNA-Seq data from other conditions become available, this list is likely to grow to include more putative RNA regulators that show differential read-through under alternative conditions.

To associate a regulator to a specific growth condition, we validated homologs of two previously identified RNA *cis*-regulators; the thiamine pyrophosphate (TPP) (preceding SP_0716) and flavin mononucleotide (FMN) (preceding SP_0178) riboswitches, using qRT-PCR (Fig 5A, C), confirming their media-specific changes in transcription. In many bacteria the TPP riboswitch binds thiamine pyrophosphate and regulates thiamine biosynthesis and transport [50]. Similarly, the FMN riboswitch regulates biosynthesis and transport of riboflavin by binding to FMN [51]. While we validated that these riboswitches respond to poor media by increasing expression of their respective genes (Fig 5A, C) we suspected that this was due to depletion of each specific ligand in the poor media. Indeed, when poor media is supplemented either with thiamine or riboflavin, expression of the TPP or FMN controlled gene (SP_0716 and SP_0178 respectively) decreases by more than 3-fold (Fig 5 B, D), suggesting that the observed differences between rich and poor media can be directly attributed to the activity of these riboswitches.

**Figure 5.**
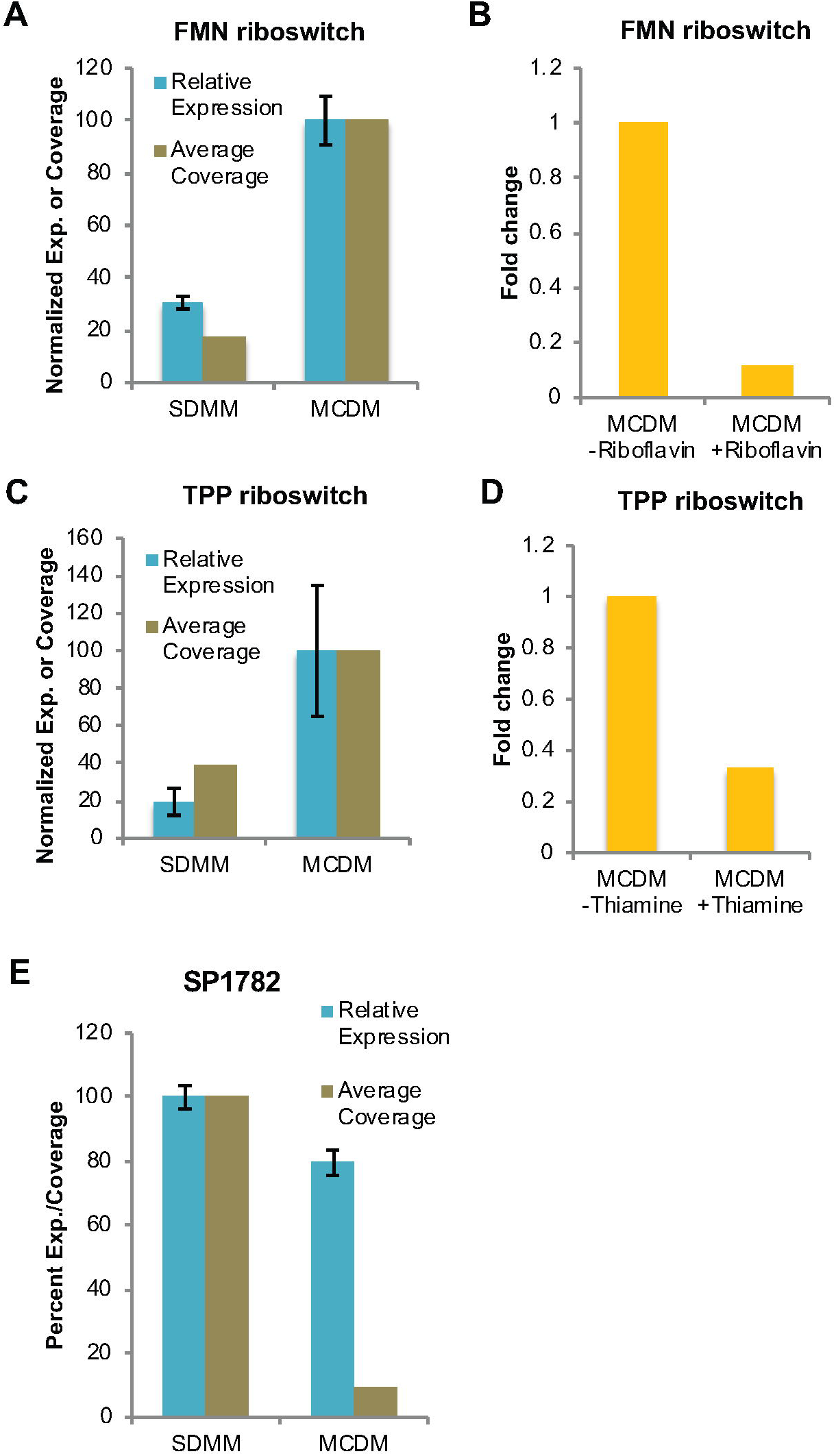
Validation of the regulatory activities of FMN, TPP riboswitches and a putative 5’-UTR regulatory candidate in different nutrient conditions. The relative expression and average RNA-Seq coverage of SP_0178 (FMN) **(A)** and SP_0716 (TPP) **(C)** increases in poor (MCDM) medium compared to rich (SDMM) medium, potentially compensating for the depletion of the specific ligand. Error bars represent standard error of the mean. Expression of SP_0178 (FMN) **(B)** and SP_0716 (TPP) **(D)** is reduced when the poor medium is supplemented with respective ligands thus confirming the regulatory activities of FMN and TPP riboswitches. (FMN-Riboflavin; TPP-Thiamine). Expression is relative to the qPCR internal control and thus for instance an expression level of 0.5 means that the expression of tested gene is 50% of the expression of internal control. Error bars represent standard error of the mean across three technical replicates. SP_1782 encoding ribosomal protein L11 methyltransferase **(E)** decreases in poor media (MCDM) compared to rich media (SDMM).

In addition to previously well-known RNA regulators, a putative regulator preceding ribosomal protein L11 methyltransferase (SP1782) that showed a decrease in read-through and expression in poor media was validated (Fig 5E). While the magnitude of the expression change apparent from qPCR is not as strong as that calculated from the RNA-Seq coverage, there are significant changes in expression that matches the direction of change. Since many ribosomal proteins are regulated through the action of cis-regulatory RNAs [52], this seems like a strong candidate for further study. However, due to the considerable challenges associated with identifying the exact ligand for this putative regulator, a definitive biological function has yet to be assigned and is currently under investigation.

In an attempt to validate the feasibility of directly associating RNA-regulators with a highly specific change in the environment we performed RNA-Seq in the presence and absence of uracil. One specific regulatory element that is sensitive to uracil is the pyrR RNA element, which in many bacteria regulates *de novo* pyrimidine nucleotide biosynthesis through a transcription attenuation mechanism mediated by the PyrR regulatory protein [53,54]. In the presence of the co-regulator UMP, PyrR binds to the 5’ UTR of the *pyr* mRNA transcript (the pyrR RNA element) and disrupts the anti-terminator stem-loop thereby promoting the formation of a factor-independent transcription terminator resulting in reduced expression of downstream genes [53] (Fig 6A). In contrast, the co-regulator 5-phosphoribosyl-1-pyrophosphate (PRPP) antagonizes the action of UMP on termination by binding to the PyrR protein when UMP concentration is low [55] (Fig 6A). For *S. pneumoniae*, our data confirms that pyrR RNA elements are present in the 5’ UTR of two *pyr* operons (SP_1278-1276; SP_0701-0702), and the uracil transporter (SP_1286). Furthermore, in response to the absence of uracil the coverage across the two genes directly adjacent to the regulators (SP_1278 and SP_0701) and over the entire two operons increases drastically (Fig 6B), demonstrating that the regulatory elements effectively turn the genes/operons on, which we confirmed by qRT-PCR (Fig 6C). Thus, while term-seq can be used to map novel regulatory RNA candidates on a genome-wide scale, RNA-Seq data can be leveraged, even in retrospect, to identify environmental conditions the regulator responds to.

**Figure 6.**
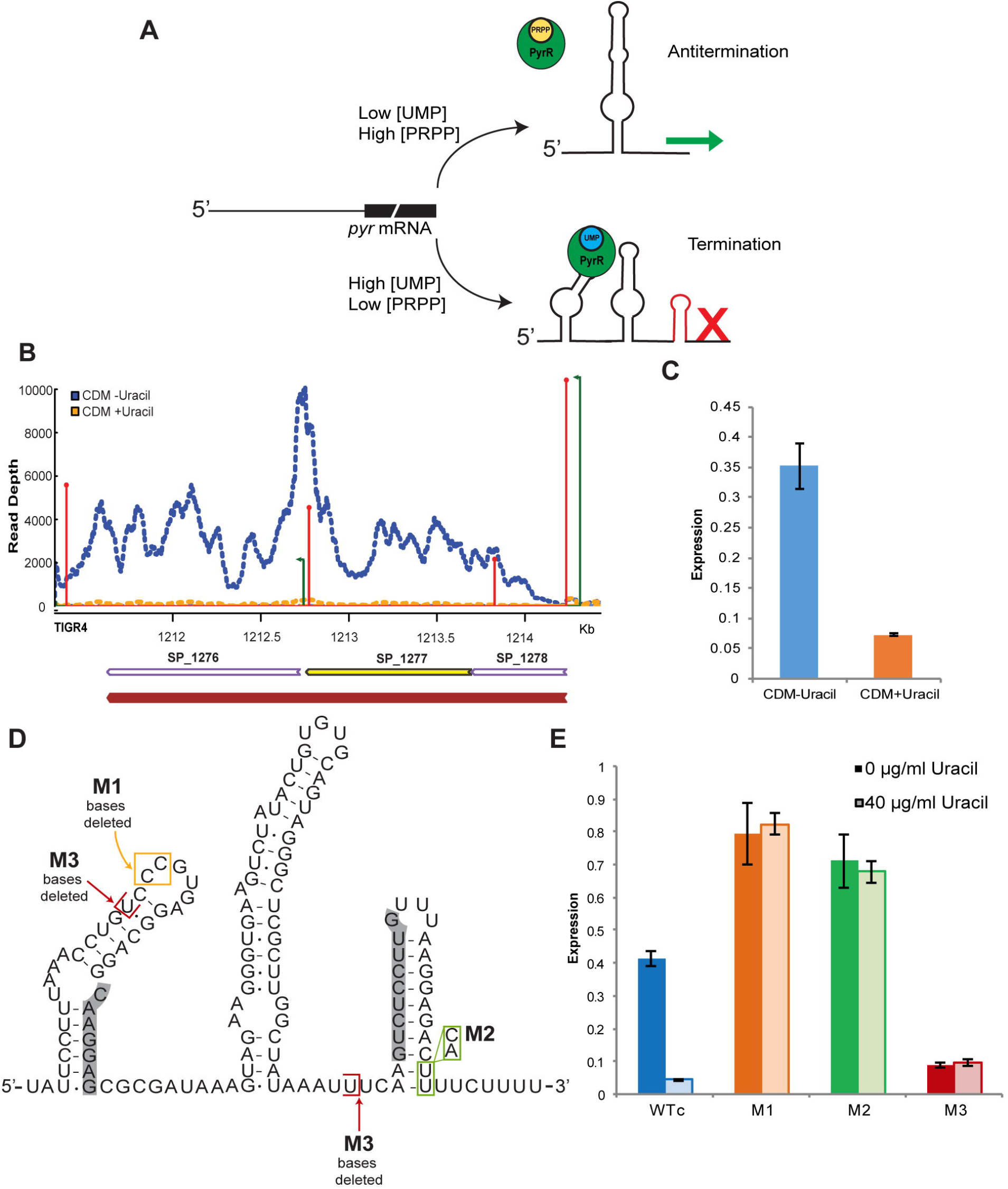
Mechanism, structure and regulatory activity of pyrR regulatory RNA element and its mutants in the presence and absence of uracil. (**A)** Schematic representation of the proposed mechanism of regulation of *pyr* operon by pyrR RNA element. In the presence of UMP, PyrR binds to the pyrR RNA and results in the formation of a premature terminator, disrupting the anti-terminator formed when UMP is low, resulting in transcription termination. **(B)** RNA-Seq coverage map across the *pyr* operon (SP_1278-1276) showing premature transcription termination and consequently decreased expression of its genes downstream of the pyrR regulator when grown in defined medium (CDM) in the presence of uracil (yellow) compared to the absence of uracil (blue). Coverage was normalized to the number of uniquely mapped reads in each library. TSSs are in green and the size represents the log transformation of the processed/unprocessed ratio and TTSs are in red and size represents the log transformation of the coverage. **(C)** qRT-PCR determining the expression of the first genes in the *pyr* operon (SP_1278) in the presence and absence of uracil validates the RNA-Seq observation. Expression is relative to the qPCR internal control. (**D)** Secondary structure of the *S. pneumoniae* pyrR RNA regulatory element in ‘off’ conformation. Boxed in red and yellow are bases that were deleted in M1 and M3 mutations, respectively. Bases boxed in green were replaced by indicated bases to make mutation M2. Highlighted in grey are nucleotides that would base pair to form the anti-terminator when the riboswitch is in the “on” conformation. (**E)** A representative qRT-PCR quantification of the expression of SP_1278 transcript from pyrR RNA mutant strains cultured in defined medium with or without uracil, corresponding to the regulatory activity of pyrR RNA mutants. While the WTc (wild type with chloramphenicol resistance cassette) decreases expression in the presence of uracil, M1 is insensitive to the ligand. M2 and M3 result in either constitutive or reduced expression of the *pyr* operon. ~2 fold higher expression in M1 compared to WTc in the absence of uracil could be the result of endogenous uracil having a slight inhibiting effect on the wild type. Expression is relative to the qPCR internal control. Error bars represent standard error of the mean across three technical replicates.

### The pyr operon is regulated through the secondary structure of the 5’ RNA leader-region, is essential for in vitro growth and in vivo virulence and can be directly manipulated

To further investigate the importance of the pyrR regulatory RNA element in growth, three different mutants were constructed that variably affect the 5’ RNA secondary structure (Fig 6D): 1) mutation M1 interferes with the binding of PyrR to the *pyr* mRNA; 2) mutation M2 renders the regulatory element in an “always on” state by destabilizing the rho-independent terminator stem-loop structure that is formed in the presence of UMP; 3) M3 locks the terminator and creates an “always off” state (Fig 6D). Wild type and mutant strains were cultured in the presence or absence of uracil and the effect of the mutations on expression of SP_1278 were assessed with qRT-PCR (Fig 6E) and further confirmed by β-galactosidase reporter assays (S4 Fig). As expected, expression in the wild type decreased (9.5-fold) in the presence of uracil confirming the repressive effect of exogenous pyrimidine (Fig 6E) [56]. M1, which should be insensitive to the presence of PyrR and its co-regulator UMP (Fig 6D) is indeed unresponsive to the presence of uracil (Fig 6E). M2 triggers constitutive expression of the *pyr* operon (Fig 6E) and M3 has a ~5-fold reduction in expression compared to the wild type regardless of the presence of uracil (Fig 6E).

Previously we showed that the pyrimidine synthesis pathway in *S. pneumoniae* is partially regulated by a two-component system (SP_2192-2193) and that genes in this pathway are important for growth [57]. To determine the importance of a functional pyrR regulatory RNA element in growth, we performed growth experiments with mutants M1, M2 and M3 in the absence and presence of uracil. These data suggest that a functional pyrR does not appear to be absolutely necessary. For instance, while M1 may have a slight growth defect when cultured in the absence of uracil, M2 has no growth defect in the presence or absence of uracil (Fig 7A, B). Although both mutations result in constitutive expression of the *pyr* operon, mutation M1 leads to higher expression (Fig 6E, S4 Fig) indicating that overexpression of the pyr genes may result in accumulation of end products that are detrimental to the cell. Alternatively, the M2 pyrR RNA element can still bind excess UMP-bound PyrR (as its PyrR binding domain is intact) thus reducing the effective concentration of UMP in the cell and thereby potential accumulation associated side-effects. Importantly, M3 has a severe growth defect compared to wild type in the absence of uracil (Fig 7A) (*p<0.002*), which can be partially rescued upon addition of uracil (Fig 7B). This suggests that while a constitutive off-state is detrimental for the bacterium in the absence of uracil a constitutive on-state can be overcome, indicating that efficient transcriptional control may not be essential.

**Figure 7.**
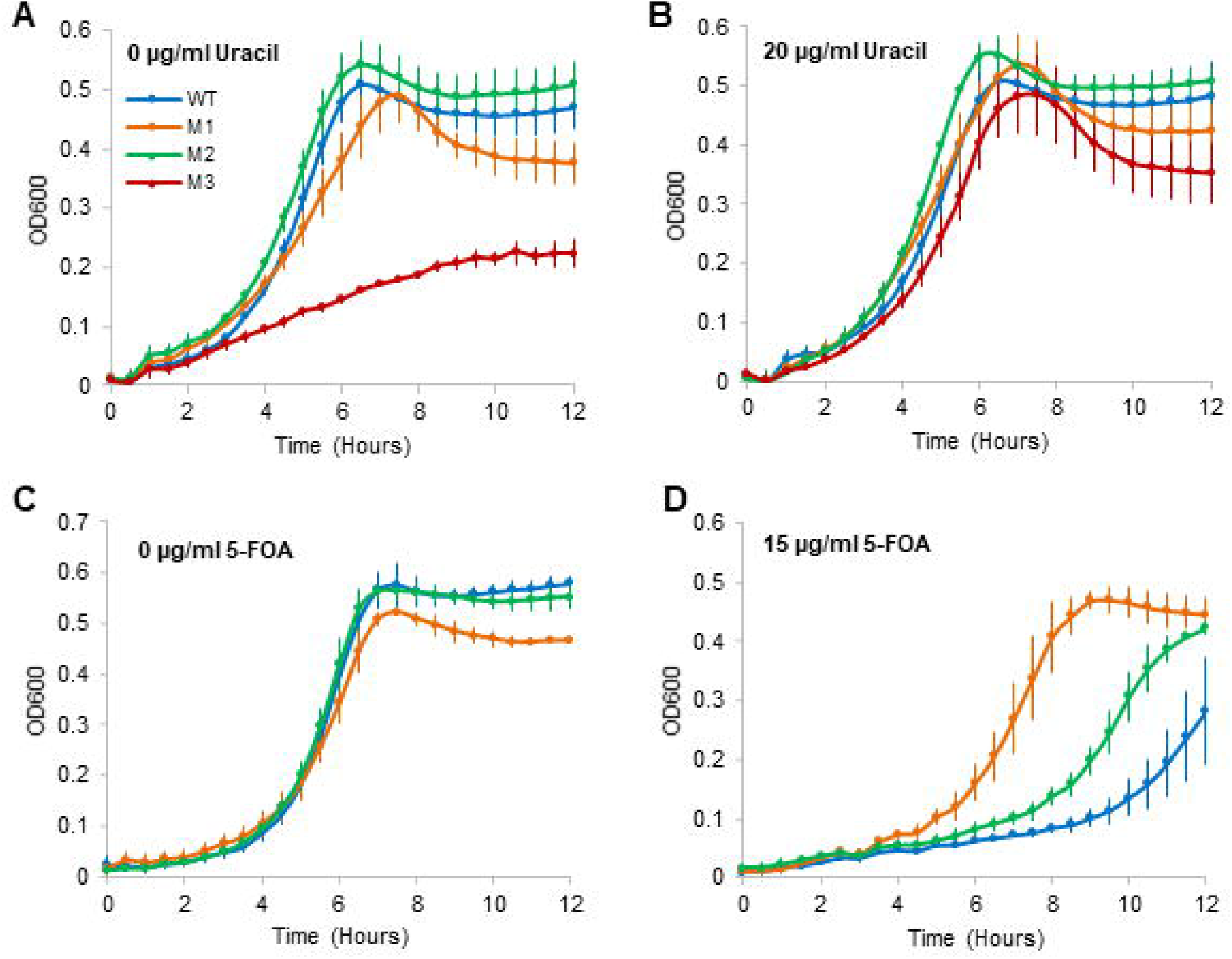
Regulation by pyrR regulatory element is important, but not essential for *in vitro* growth of *S. pneumoniae*. **(A-B)** *In vitro* growth curves of mutants when cultured in defined media with (20 μg/ml) or without uracil. While mutant M2 (green) does not display a growth defect, mutants M1 (orange) and M3 (maroon) have growth defects that are restored in the presence of uracil indicating that a functional pyrR RNA element is important, but not absolutely essential for *in vitro* growth of *S. pneumoniae*. WT (blue) does not have a growth defect in the tested conditions. **(C-D)** Representative *in vitro* growth curves of mutants cultured in media with (15 μg/ml) or without 5-FOA, a toxic uracil analog. All strains show varying degrees of defects in the tested conditions indicating that a drug targeted against the secondary structure can severely and specifically hamper growth. Mutant M3 was not included in this assay as it cannot be grown without uracil. Error bars represent standard error of the mean across three biological replicates.

To determine whether we can manipulate the manner in which the pyrR RNA element affects growth, we determined growth in the presence of 5-Fluoroorotic acid (5-FOA), a pyrimidine analog. 5-FOA is converted into 5-Fluorouracil (5-FU) a potent inhibitor of thymidylate synthetase, whose activity is essential for DNA replication and repair [58]. Additionally, 5-FU competes with UMP for interacting with the PyrR protein [59]. 5-FU can thus work as a decoy, signaling that UMP is present in the cell; triggering the formation of a terminator and reducing expression of the *pyr* operon. The wild type strain displayed a severe growth defect in the presence of 5-FOA (Fig 7C&D) (*p<0.05*), while M1 (which should not interact with PyrR and should thus be largely insensitive to the presence of 5-FOA) displayed a much smaller growth defect (Fig 7C&D). In addition, M2, which constitutively over expresses the *pyr* operon, is also less sensitive to 5-FOA then wild type (Fig 7C&D) (*p<0.05*). Thus, the mutations we introduced into the pyrR RNA element affect the secondary structure in the manner that we intended, and can have far reaching regulatory and fitness effects. Importantly, it shows that a drug targeted against the secondary structure can directly manipulate and severely hamper growth.

A remaining key question is the importance of RNA regulatory elements in colonization and the induction of disease. Somewhat surprisingly our *in vitro* growth curves suggest that constitutive expression of the *pyr* operon (M1) and constitutive overexpression (M2) is not substantially detrimental to growth, indicating that efficient regulation is not critical. To assess the effect of loss of regulation on bacterial fitness *in vivo*, the pyrR RNA element mutants were tested in 1×1 competition assays (mutant vs. wild type) in a mouse infection model (Fig 8). While fitness for all three mutants is similar to the unmodified strain *in vitro* in the presence of uracil, M1 and M3 are unable to colonize and survive in the mouse nasopharynx, or infect and survive in the lung and transition and survive in the blood (Fig 8A&C). M2 has less of a defect *in vivo*, but still has a significantly diminished ability to infect and survive in the lung (Fig 8B). These results indicate that efficient regulation of the *pyr* operon *in vivo* is critical for growth and survival of *S. pneumoniae* within the host. While we had previously shown that genes in the *pyr* operon are important *in vivo* [57], the regulatory findings in this project take our understanding a step further and, importantly, in combination with the findings that 5-FOA can efficiently interact with the RNA regulatory element, suggests that it is feasible to modulate *in vivo* fitness and thereby virulence by targeting such regulatory elements.

**Figure 8.**
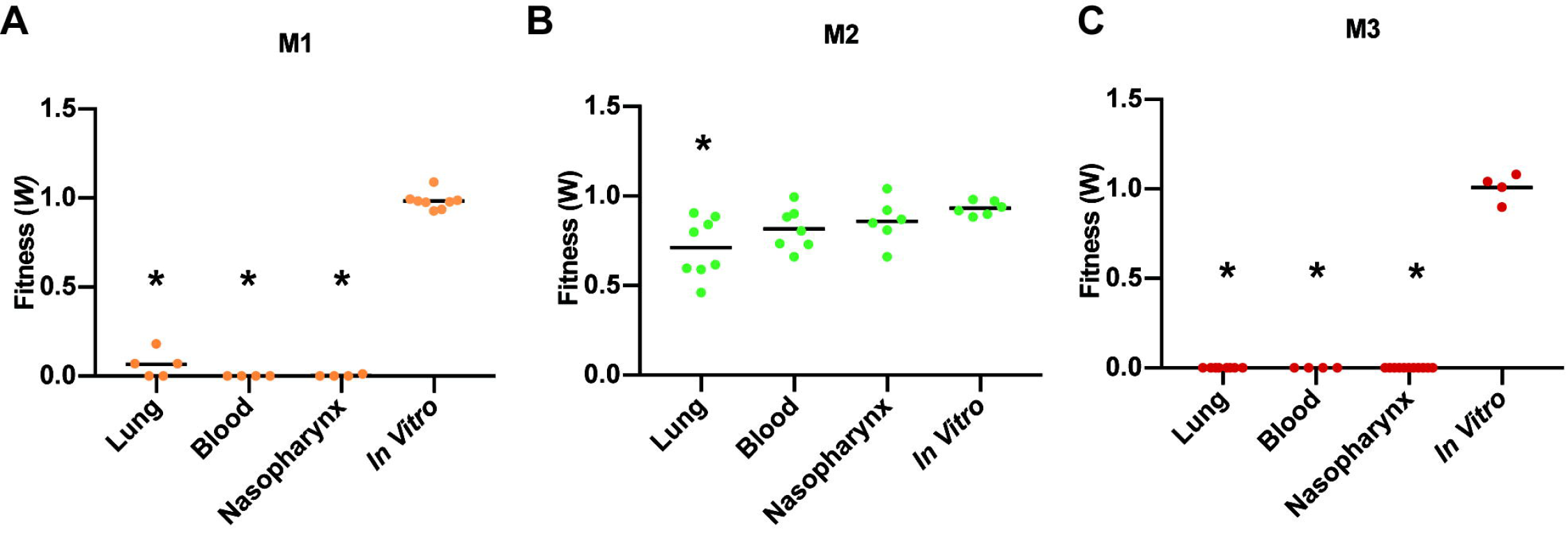
Regulation by the pyrR RNA element is crucial for *in vivo* survival and virulence of *S. pneumoniae*. 1×1 competition assays (mutant vs wild type T4) reveals fitness defects of pyrR RNA mutants in a mouse infection model. While M1 (A) and M3 (C) display severe defects in all the tested *in vivo* environments namely lung, blood and nasopharynx, M2 (B) has less of a defect. Significant change in fitness (*p<0.0125)* are indicated by asterisks (*). Each data point represents a single mouse.

### Towards comprehensive transcriptional landscape reconstructions and highly targeted regulatory RNA element inhibitors

With the advent of deep sequencing technologies, our understanding of prokaryotic transcriptional dynamics is rapidly advancing [60] and underlining that bacterial transcriptomes are not as simple as previously thought. Analysis of the *S. pneumoniae* TIGR4 transcriptome using three different sequencing techniques (RNA-Seq, term-seq, and 5’end-Seq) has led to a comprehensive mapping of its transcriptional landscape. We successfully map 742 primary and 48 secondary high confidence TSSs and 74 low confidence secondary TSSs (2150 total) and 1864 TTSs. These results along with a recent study that annotates the *S. pneumoniae* D39 genome [61], deeply enriches the understanding of transcriptional regulation in this important pathogen. Besides identifying the transcript boundaries, we also uncovered a complex operon structure, which has also been found in *E. coli* [4]. Importantly, such complexity likely allows for environment-dependent modulation of gene expression producing variable transcripts in response to varying conditions, which we illustrated here through analyses of a 9-gene complex operon and the mal regulon (Fig 4). Additionally, similar environment-dependent versatile operon behavior has been observed in *E. coli* [4] and to a lesser extent in *Mycoplasma pneumoniae* [62]. This means that our understanding is shifting dramatically and it is thus becoming clear that operons in bacteria should be seen as adaptable structures that can significantly increase the regulatory capacity of the transcriptome by responding to environmental changes in a highly specific manner. Furthermore, this technology can potentially be extended to *in vivo* growth environments to determine how the bacteria modulates its gene expression in the context of the host immune system. A major challenge with respect to such procedures is the quantity and quality of RNA to generate TSSs, TTSs and RNA-Seq coverage data. However, such limitations could be overcome by obtaining only RNA-Seq coverage data and leveraging these data, as we have done in this study, to identify RNA regulators specific to *in vivo* environments.

Another central aspect of our approach is the identification of putative 5’-UTR structured regulatory elements. Riboswitches and other untranslated regulatory elements (binding sites for small regulatory RNAs) are important bacterial RNA elements that are thought to regulate up to 2% of bacterial genes [7,8]. However, the discovery of new regulators is difficult when relying solely on computational methodology and sequence conservation [63]. Here we show that through term-seq [35] it is possible to identify such RNA elements on a genome-wide scale and by combining it with RNA-Seq performed in different conditions transcriptional phenotypes can be directly linked to the RNA element. This strategy thus makes it possible to screen for regulatory RNA elements in retrospect by making use of already existing or newly generated RNA-Seq data.

Importantly, besides the ability to re-construct an organism’s intricate transcriptional landscape we show that there is also a direct application of our multi-sequencing approach, namely the ability to inhibit operons and/or pathways with specific chemicals or drugs that target the RNA regulatory element. We show that this is possible for the pyrR RNA element, a regulatory element that is important for pneumococcal growth and virulence, which means that this regulatory element could be a potential antimicrobial drug target. This idea is further strengthened by the fact that *S. pneumoniae* displays a growth defect in the presence of 5-FOA, which directly relates to misregulation of pyrR RNA confirming its drug-able potential.

We believe that the presented multi-omics sequencing strategy brings a global understanding of regulation in *S. pneumoniae* significantly closer, and because the approach is easily transferable to other species, it will enable species-wide comparisons for conservation of operon structure and regulatory elements. In addition, such detailed regulatory understanding creates new regulatory control tools for synthetic biology purposes. Moreover, the combination with *in vivo* experiments shows that it is a realistic goal to design or select specific compounds that target ribo-regulators in order to mitigate virulence or antibiotic resistance.

## Methods

### Culture conditions and sample collection

For RNA-Seq, term-seq and 5’end-Seq library preparation, *Streptococcus pneumoniae* TIGR4 (T4) was cultured in rich media (SDMM) to mid-log phase (OD_600_ = 0.4). Cultures were diluted to an OD_600_ of 0.05 in fresh media, grown for one doubling (T_0_). Cultures were then grown in the presence or absence of 0.24 μg/ml vancomycin. At 0 min (T_0_) and after 30 min of growth (T_30_) 10ml culture was harvested by means of centrifugation (4000 rpm, 7 min at 4°C) followed by flash freezing in a dry-ice ethanol bath and storage at −80°C until RNA extraction. Sample collection was performed in four biological replicates and total RNA was isolated using an RNeasy Mini kit (Qiagen). For qRT-PCR analyses, T4 was cultured in SDMM to mid-log phase (OD_600_ = 0.4) and after centrifugation cultures were washed with 1X PBS and diluted to an OD_600_ of 0.003 in appropriate media. Cultures were harvested at mid-log followed by RNA extraction as described above.

### 5’end-Seq library preparation

5’end-Seq libraries were generated by dividing the total RNA into 5’ polyphosphate treated (processed) and untreated (unprocessed) samples that were subsequently processed and sequenced according to protocols described in [22] and [64] with few modifications. See supplemental methods for a detailed protocol.

### RNA-Seq library preparation

RNA-Seq libraries were generated by using the RNAtag-Seq protocol [34,44]. Briefly, 400 ng RNA was fragmented in FastAP buffer, DNase-treated with Turbo DNase, 5’-dephosphorylated using FastAP. Barcoded RNA adapters were then ligated to the 3’ terminus, samples from different conditions were pooled and ribosomal RNA was depleted using the Ribo-zero rRNA removal kit. Illumina cDNA sequencing libraries were generated by first-strand cDNA synthesis, 3’ linker ligation and PCR with Illumina index primers for 17 cycles. The final concentration and size distribution were determined with the Qubit dsDNA BR Assay kit and the dsDNA D1000 Tapestation kit, respectively.

### Paired-end RNA-seq library preparation

RNA was extracted in triplicate from cells grown to mid-logarithmic phase (OD_620_=0.4) using the QiaShredder and RNeasy Kit (Qiagen) by following the kit instructions with the exception of the lysis step, which was performed by membrane disruption using zirconia/silica beads. After extraction, RNA was treated with DNA Free DNA removal kit (Thermo) to remove DNA. RNA-seq library was generated via ScriptSeq vs RNA-seq library kit (epicenter) using 1 ug RNA. The kit instructions were followed using Ampure beads (Agencourt) for cDNA and library purification. Each replicate was individually indexed using Failsafe PCR enzyme (Epicentre). Completed libraries were analyzed for insert size distribution on a 2100 BioAnalyzer High Sensitivity kit (Agilent Technologies, Santa Clara, California). Libraries were quantified using the Quant-iT PicoGreen ds DNA assay (Life Technologies,). One hundred cycle paired end sequencing was performed on an Illumina HiSeq 4000.

### term-seq library preparation

term-seq libraries were generated as previously described [35] with few modifications. 2 μg total RNA was depleted of genomic DNA using Turbo DNase, 5’ dephosphorylated, ligated to barcoded RNA adapters at the 3’ terminus and fragmented in fragmentation buffer. Barcoded and fragmented RNA from different conditions were pooled and ribosomal RNA was depleted using Ribo-zero. cDNA libraries were generated by first strand cDNA synthesis and RNA template was degraded as mentioned in the 5’end-Seq library preparation. Second 3’ linker was ligated and PCR amplified with Illumina index primers for 17 cycles. All four library preparations (RNAtag-Seq, term-seq, 5’end-Seq processed and 5’end-Seq unprocessed) were pooled according to the method of preparation and sequenced at high depth (8.5 million reads/sample) on an Illumina NextSeq500.

### Read processing and mapping

The *S. pneumoniae* TIGR4 (NC_003028.3) genome annotation was supplemented with all known structured ncRNAs and sRNAs. Homologs of known ncRNA families deposited in Rfam [65,66] were identified in the genome by the cmsearch function of Infernal 1.1 [36]. sRNAs previously identified in *S. pneumoniae* [16,18] were identified in T4 using BLAST [67]. Coordinates for the hits from the above searches are included in the supplemented annotation. The single-end sequencing reads from the 5’ end-Seq sequencing, 3’ end sequencing (term-seq), and RNAtag-Seq were processed and mapped to the supplemented *S. pneumoniae* TIGR4 (NC_003028.3) genome using the in-house developed Aerobio pipeline. Aerobio runs the processing and mapping in two phases. Phase 0 employs bcl2fastq to convert BCL to fastq files, quality control and de-multiplexing and compilation of the reads based on the sample conditions. Phase 1 maps the de-multiplexed reads against the genome, under default parameters, using Bowtie2 [68] and streams the uniquely mapped reads to SAMtools [69] to generate sorted and indexed BAM files for each sample.

### in silico prediction of transcription start sites (TSSs) and transcription termination sites (TTSs)

Perl code from [35] was adapted to map the strand specific single nucleotide coverage of the 5’ positions of the 5’ end and 3’ end sequencing reads respectively estimated using BEDTools [70]. With this nucleotide level coverage data calculated from the 5’ end-Seq, regions up to 500 nucleotides upstream of the translational start sites described in the annotated TIGR4 genome (NC_003028.3) were scanned for mapped reads with a minimum coverage of 2 and a processed/unprocessed ratio of 1 as in [35]. Predicted primary TSSs were called as high confidence if the position had a minimum processed coverage greater than 100 and a processed/unprocessed ratio of 4. Secondary TSSs are greater than 10 nucleotides up or downstream of predicted primary TSSs. When multiple putative TSSs were identified in a 5’ UTR, the higher confidence, or the highest processed/unprocessed ratio was assigned as the primary TSS for the downstream gene.

Regions up to 100 nucleotides upstream of the predicted high confidence TSSs were split into three datasets – TSS-20; TSS-20-50; and TSS-100, and searched for enriched promoter motifs and transcription factor binding sites listed in [71] using MEME [72]. Similar to the identification of the TSSs, TTSs were identified by scanning up to 150 nucleotides downstream of the translational stop site for mapped 3’ end reads with a minimum coverage of 10 in the pooled dataset. The position with the highest coverage was considered the most likely TTS for a gene. Predicted TTSs were called as high confidence if the coverage of the site was at least double the average coverage in the 150 nucleotide region. 40 nucleotides upstream of the predicted TTS were scanned for a stable stem-loop structure using RNAFold [73], and the number of uridines in the 8 nucleotides immediately upstream calculated. The modified perl scripts, sample input files, and the track files for visualization are available at https://github.com/nikhilram/T4pipeline.

### Identification of transcript boundaries and operon structures in the genome

Paired-end reads were mapped to the T4 genome using TopHat2 [74]. The BAM file of the mapped reads were quality filtered and used to predict transcript boundaries and group genes into operons using a stringent run of StringTie [37,38]. Specifically, 0 locus gap separation was allowed and a minimum junction coverage of 10 was required. Predicted operons were compared with the genome based predictions listed in the Database of Prokaryotic Operons [5,42,75], and complexity in the operon structure was characterized by surveying the number of high confidence primary internal TSSs and enriched TTSs similar to [4].

### Identification of candidate regulatory elements

Once the TSSs were identified, 5’-UTR regions with a length of at least 70 nucleotides were scanned for mapped 3’ end sequencing reads with a minimum coverage of 2 to identify putative early terminators. 5’-UTR regions with a predicted early TTS were binned as candidate regulatory elements. The nucleotide sequence for each candidate element was obtained and folded using RNAFold [73]. Secondary structures and free energy values were compiled for each candidate.

The response of candidate regulatory elements to different media conditions were assessed by calculating the RNA-Seq coverage in both the regulatory element and the regulated gene. Read-through was calculated for each of the candidates as described previously [35]. Briefly, read-through is the ratio (denoted in percentage) of the average coverage across the gene to that of the 5’-UTR identified here. The greater the read-through, the higher the expression of the gene with respect to the 5’-UTR. That is, if the regulator reduced the expression of the gene, read-through would be small. If the regulator turned on gene expression in response to certain conditions, the read-through would be large.

### Conservation of the candidate regulatory elements in Streptococcus pneumoniae

A local BLAST [67] database was generated with the genomes of 30 *S. pneumoniae* strains available in Refseq 77 [76] and 350 strains from [48]. Each of the candidate regulators identified in the genome of TIGR4 was BLASTed against this database, and hits in the other genomes were extracted and aligned using MAFFT version 7 [77]. The degree of conservation across the 380 genomes was determined by surveying each candidate cluster post filtering to remove sequences that were less than 70% in length of the query and with e-values greater than 1×10^-4^.

### Expression analysis using qPCR

RNA was isolated from cultures using the Qiagen RNeasy kit (Qiagen). DNase treated RNA was used to generate cDNA with iScript reverse transcriptase supermix for RT-qPCR (BioRad). Quantitative PCR was performed using a Bio-Rad MyiQ. Each sample was normalized against the 50S ribosomal gene, SP_2204 and were measured in biological replicate and technical triplicates. No-reverse transcriptase and no-template controls were included for all samples. Primers used in this study are listed in Table S7.

### pyrR RNA mutant strain construction

To generate a pyrR RNA mutant strain, wild-type pyrR RNA element region and ~ 1 kb regions of homology flanking on either side of the RNA element was PCR amplified from *S. pneumoniae* TIGR4 genomic DNA (GenBank accession number NC_003028.3). Mutations to the pyrR RNA element were obtained either by site-directed mutagenesis or by PCR assembly using appropriate primers (S7 Table). A chloramphenicol resistance cassette was PCR amplified from pAC1000. The amplified products were assembled such that the chloramphenicol resistance cassette was inserted immediately upstream of the promoter in the opposite orientation. The assembled products were transformed into *S. pneumoniae* TIGR4 as previously described [78] and transformants were screened for resistance to chloramphenicol and mutation verified by Sanger sequencing.

### pyrR RNA mutant growth assays

Wild type and pyrR RNA mutants of T4 were grown for 2 hours and diluted to an OD_600_ of 0.015 in fresh media, with varying concentrations of uracil and/or 5-FOA. Growth assays were performed in 96-well plates for 16 hours by taking OD600 measurements every half hour using a Tecan Infiniti Pro plate reader (Tecan). Growth assays were performed no less than three times.

### In vivo pyrR mutant fitness determination

1 x 1 competition experiments were performed with pyrR RNA mutants (M1 to M3) that were competed against the wild-type strain after which bacterial fitness was calculated as previously described [57] with a few modifications. Lung removal and homogenization (in 10 ml 1 x PBS), blood collection (100 μl) and nasopharnyx lavage (with 1 ml 1X PBS) were perfromed on all animals 24 hours post infection, with the exception of pyrR M3, which due to the large fitness defect were harvested at 6 hours post infection. Experiments involving animals were performed in accordance with guidelines of IRB/IACUC and approval of the Boston College Animal Facility.

## Acknowledgements

We would like to thank Jon Anthony for sequence data processing on the Aerobio platform, Andrew Goodwin for technical assistance with qPCR, Charles S. Hoffman for the generous gift of 5-FOA and Daniel Dar for discussion and helpful suggestions. We would also like to thank Jelle Slager and Jan-Willem Veening for their in-depth critique of the manuscript. The sequencing datasets generated during the current study are available in the Sequence Read Archive (SRP136114).

---

**Supplemental figure 1. Confidence in the predicted TSSs. A.** Frequency distribution of the distance between the TSS position predicted for each gene between conditions. This also takes into account whether TSSs were predicted. Majority of the predictions match exactly or are very close to one another between the conditions providing confidence in the TSS predictions from the pooled data. **B.** Predicted TSSs show enrichment of the TATA box at the −10 and the TTGACA at the −35 positions respectively as identified by MEME [72].

**Supplemental figure 2. Differences in the operon predictions between StringTie and Rockhopper. (A)** StringTie reconstructed transcript grouping genes SP_0001 – SP_0014. Log transformed paired-end RNA-Seq coverage map across this region. Red and Green lines represent the predicted TSSs and TTSs **(B)** The pie chart describes the distribution of the types of operons present in T4 identified using single-end RNA-Seq analyzed with Rockhopper. A total of 474 multigene and 773 single gene operons were identified, which can be divided up in 63% simple operons (single gene transcriptional units with a single TSS and TTS; gray), 29% complex operons (multi-gene operons with multiple TSSs and TTSs; orange), 4% traditional operons (multi-gene operon with a single TSS and TTS; green), 1% multiTSS operons (blue), and 3% multiTTS operons (red). (**C)** The clustered histogram describes the distribution of genes and transcriptional features in non-traditional operons described by Rockhopper, where gray represents the numbers of genes in the multigene operons, green represents the number of TSSs within operons, red represents the number of TTSs within operons. Two-gene operons are found most frequently in the non-traditional operons with one internal TSS and TTS.

**Supplemental figure 3. Distribution and conservation of the 141 putative regulatory candidates across 380 *S. pneumoniae* genomes. A.** Frequency distribution of the candidates across the surveyed genomes. **B.** Conservation of the candidates as a measure of the mean p-Distance within each candidate cluster.

**Supplemental figure 4. β-galactosidase activities of pyrR mutant strains in the presence or absence of uracil.** The activity of the wild type pyrR decreases in the presence of uracil, while M1 is insensitive to the ligand. Activity of mutant M2 results in increased Miller units even in the presence of uracil. Empty *lacZ* represents a β-galactosidase reporter construct without a regulatory region upstream, which was used as a negative control. Error bars represent standard error of the mean across three technical replicates.

**Supplemental table 1. List of all the transcriptional start sites (TSSs) identified from the 5’ end-Seq in *S. pneumoniae* strain TIGR4.** The 742 high confidence primary TSSs (sheet 1), the 48 putative secondary high confidence TSSs (sheet 2), 1286 low confidence primary TSSs identified (sheet 3), and the 74 low confidence secondary TSSs (sheet 4), their Processed/unprocessed ratios and coverages, and the lengths of the 5’-UTR thus formed are listed. Additionally, a description of whether the TSS is intergenic or within an upstream coding sequence is provided.

**Supplemental table 2 List of all the transcription termination sites (TTSs) identified from term-seq in *S. pneumoniae* strain TIGR4.** The positions of all 1864 enriched sense TTSs, and the coverage of each site pooled from all the samples are listed. Lengths of the 3’-UTRs are also listed for the sense TTSs. The predicted stem-loop structure and the number of uridines upstream of each TTS is also shown.

**Supplemental table 3. All the multi-gene operons identified by StringTie.** A description of the 388 multi-gene operons identified by StringTie. Number and the locus tags of genes that are in operons.

**Supplemental table 4. Table of candidate 5’-UTR regulatory elements.** Detailed description of the candidate 5’-UTR regulatory elements for 141 genes. Position of the TSS and its Processed/unprocessed ratio, early TTS and average coverage for each are described. Additionally, length of the 5’-UTR as well as whether it is intergenic or within the upstream coding sequence is provided.

**Supplemental Table 5. RNA adapters with barcode for 5’ end-Seq.** List of adapters used for 5’ end-Seq. Highlighted in red are the barcode sequence (5’-3’).

**Supplemental Table 6. Primers used in the study.**

